# MacAction: Realistic 3D macaque body animation based on multi-camera markerless motion capture

**DOI:** 10.1101/2024.01.29.577734

**Authors:** Lucas M. Martini, Anna Bognár, Rufin Vogels, Martin A. Giese

**Affiliations:** Hertie Institute for Clinical Brain Research (HIH) and Centre for Integrative Neuroscience (CIN), University Clinic Tübingen, Tübingen, Germany; International Max Planck Research School for Intelligent Systems (IMPRS-IS), Tübingen, Germany; Department of Neuroscience, KU Leuven, Leuven, Belgium; Leuven Brain Institute, KU Leuven, Leuven, Belgium

**Author notes:** Contributing authors.

## Abstract

Social interaction is crucial for survival in primates. For the study of social vision in monkeys, highly controllable macaque face avatars have recently been developed, while body avatars with realistic motion do not yet exist. Addressing this gap, we developed a pipeline for three-dimensional motion tracking based on synchronized multi-view video recordings, achieving sufficient accuracy for life-like full-body animation. By exploiting data-driven pose estimation models, we track the complete time course of individual actions using a minimal set of hand-labeled keyframes. Our approach tracks single actions more accurately than existing pose estimation pipelines for behavioral tracking of non-human primates, requiring less data and fewer cameras. This efficiency is also confirmed for a state-of-the-art human benchmark dataset. A behavioral experiment with real macaque monkeys demonstrates that animals perceive the generated animations as similar to genuine videos, and establishes an uncanny valley effect for bodies in monkeys.

## Introduction

Understanding the computational neural processes in social vision remains an unresolved problem in neuroscience. Because primates are social animals, evolution has equipped their brains with special neural mechanisms for facial and body recognition [1]. The clarification of these mechanisms necessitates experiments with highly controlled visual stimuli that manipulate underlying visual features [2, 3]. For the study of social perception, avatars, as opposed to real-world videos of behaving animals, offer the advantage of precise control of relevant stimulus features and timing. To justify a transfer of the obtained results to real stimuli, such avatars need to exhibit sufficient realism in both shape and motion.

While there has been notable progress in generating dynamic macaque face avatars [4, 5], there have so far been no attempts to create lifelike dynamic monkey body avatars with realistic motion. This gap can be attributed, in part, to the challenges associated with marker-based motion capture in macaques. Not only is the reproducible positioning of markers on their dense fur technically demanding, but these animals also typically remove or destroy body-attached markers during grooming. Consequently, various datasets have been constructed to capture poses in images [6–8]. Such tracking of non-human primates is challenging, because of the high-dimensional pose space, the presence of strong occlusions, and the homogeneity of texture induced by fur. Therefore, recent multi-camera approaches have reconstructed 3D poses only by exploiting setups with large numbers of cameras and extensive hand-labeling of keypoints [9–11]. These disadvantages render such approaches infeasible as a basis for the estimation of 3D joint angle trajectories for computer animation, due to the larger number of keypoints and high accuracy required.

In other animals with less complex pose spaces, the surface models of avatars were incorporated in the tracking process [12–14]. Others bypassed the use of commercial animation software entirely by applying neural rendering, which creates photorealistic animations directly from image data [15–17], while sacrificing the possibility to control and parameterize individual stimulus features.

Computer animation stimuli raise the question of ecological validity, and even stimuli that appear realistic to humans might not do so for monkeys. In general, when the realism of artificial agents is continuously increased, an uncanny valley might emerge, where preference for the animation decreases close to the highest levels of realism [18] (Fig. 4b). This phenomenon has been exploited to verify whether graphics methods reach a sufficient level of realism [19]. An uncanny valley has also been demonstrated for macaque observers for static [20] and dynamic monkey faces [5]. Whether such an effect also exists for monkey bodies is unknown.

Addressing these challenges, we here introduce MacAction, the first approach for the generation of highly realistic animations of macaques based on markerless video motion capture, exploiting commercial animation software. The developed pipeline processes high-resolution, multi-camera footage with state-of-the-art pose detectors, interpolating an entire action sequence by a minimum number of labeled keyframes. Thereby, our approach enables accurate tracking using large marker sets, as required for 3D animation, with a very small amount of required hand-labeling.

Achieving realistic single-action animations from 6 cameras becomes feasible with only 2 labeled frames per second (LPS), where our method substantially reduces errors compared to other macaque pose estimation methods [21]. Through a behavioral experiment with macaques, we show the existence of an uncanny valley for bodies in macaque perception. In fact, our animations (Supplementary Videos 1 and 2) elicit distributions of attention comparable to real-life video footage, making such artificial stimuli suitable for the study of social perception in monkeys.

## Results

### MacAction: a framework for 3D animation of highly realistic macaque avatars

To create realistic artificial stimuli that offer a high level of stimulus control, we devised a pipeline for the animation of a commercial macaque avatar model with 86 joints, with a total of 261 degrees of freedom (including global translation), exploiting markerless tracking. Realistic animation necessitates accurate 3D tracking of a sufficient number of keypoints in order to constrain the high-dimensional body pose (e.g., rotations of the arms, wrists, feet, finger movements, and the tail). Our work is based on 42 keypoints, a number that exceeds by factor two the one in previous studies on markerless tracking of macaques [6, 7]. For the generation of highly realistic scenes, we also reconstructed the background in computer graphics using 3D modeling software based on laser scans of the monkey cage (Fig. 1c).

**Fig. 1.**
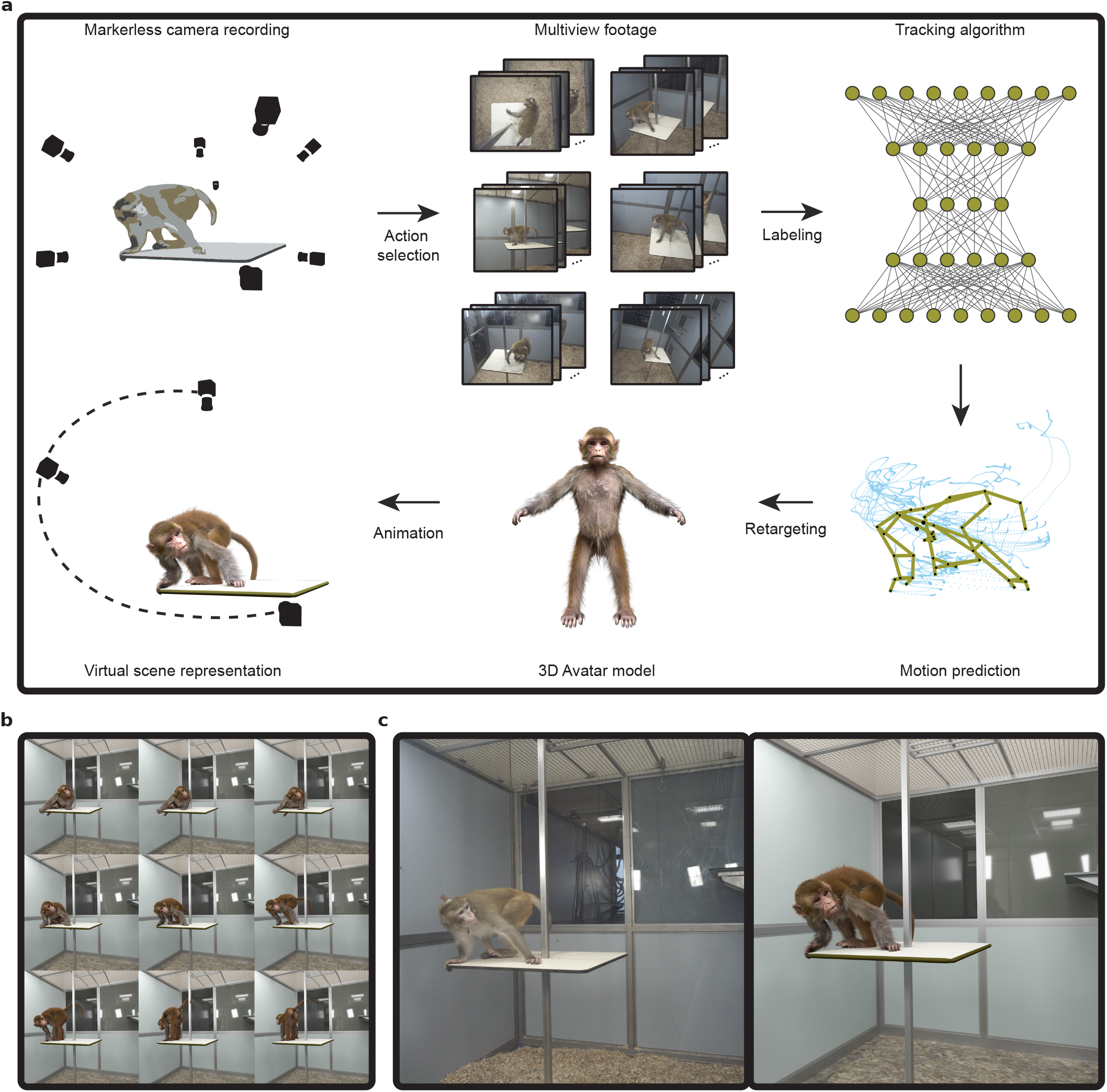
MacAction animates 3D markerless pose tracking to generate realistic synthetic images. **a**, Pipeline for character animation: Selection of actions from sets of synchronously recorded videos from multiple viewpoints. A neural network-based tracking algorithm predicts keypoint trajectories for all frames of an action (shown in blue) based on a minimal subset of hand-labeled data for training. The retargeting process then maps this motion onto a 3D avatar. The resulting animation can be embedded into a 3D model of the background scene, which can be rendered from any viewpoint. **b**, Generated images of the virtual action over time. **c**, Comparison between corresponding frames from a real video (left), and the simulated virtual scene (right), combining the animated avatar with the 3D background model.

From 150 minutes of synchronously recorded multi-view video footage, we selected three distinct actions, containing a single and multiple animals: (1) a complex submissive turning movement; (2) a short walk representing an emotionally neutral behavior; and (3) a social interaction of two individuals, where one monkey grooms the other.

State-of-the-art markerless tracking methods for animals [22–24] require human annotators to label exemplary postures for the training of underlying neural networks. For computer animation, however, the required amount of labeling would significantly escalate. Reducing manual labor is thus a key challenge in making our approach feasible for applications in neuroscience. To simplify the labeling process in multi-camera footage, we developed a software tool, AnimLabel, that propagates constraints from already annotated views to unlabeled ones.

The resulting pairs of multi-camera images and 3D poses served as training examples for state-of-the art markerless tracking algorithms, reconstructing poses for intermediate unlabeled time points. We first tested DeepLabCut (DLC) [22, 25], a popular image-based tracking algorithm for different animal species. To generate 3D poses, we processed tracked 2D keypoints from multiple views, exploiting the software Anipose [26]. This allowed us to incorporate spatiotemporally regularized triangulation that constrains 3D keypoints by optimizing their projections across views, their temporal smoothness, and minimizing the variance of their distance to neighboring markers. The two control parameters *α*_*time*_ and *α*_*limb*_ allow us to steer the importance of the latter objectives within the triangulation. We refer to the process of discarding those factors in the triangulation as structural refinement.

We also applied another approach, originally developed for human tracking [27], which integrates the triangulation process into the learning objective of the neural network. Its adoption for animals, the 3-Dimensional Aligned Neural Network for Computational Ethology (DANNCE), outperformed DLC with the same number of cameras for behavioral tracking in rats and mice [24]. The tracking results obtained with these two methods were further processed and mapped onto the skeleton of the macaque avatar. For this purpose, we used custom scripts in Maya for skeleton retargeting, making the pipeline adaptable to different marker configurations. Using smoothed 3D marker trajectories, we were able to compute the poses of the avatar for each time step and to animate the actions in a virtual scene with a sufficiently high framerate (Fig. 1a,b).

### Accurate single-action tracking with minimal labeling

Previous work has used extensive numbers of cameras and labels to realize markerless motion capture for macaques, mainly aiming at behavioral analysis [9]. However, cameras and labeling work are costly resources, posing a challenge for smaller labs. We, therefore, validated the developed method in terms of the number of labeled frames required for accurate tracking, dependent on the number of cameras and the used tracking algorithm.

We first considered the densely hand-labeled submissive action (4.38 LPS). The DLC 2D landmarks were triangulated with spatiotemporal regularization in Anipose [26], but omitting temporal and limb constraints. As the pose predictions were noisy, we initialized the triangulation by Random Sample Consensus (RANSAC). With six labeled frames, this method achieved reasonable test error performance (mean error and SD, 18.9 *±* 20.8mm), close to DANNCE’s voxel resolution. It even outperformed DANNCE when training with double the number of labeled frames (29.3 *±* 25.9mm) for the same number of cameras (Fig. 2a,b and Supplementary Fig. 1a). However, the pose trajectories obtained by DANNCE were smoother (Fig. 2d and Supplementary Videos 3 and 4). This was quantified by computing the average of the predicted marker speed across the sequence, termed instantaneous per-joint temporal deviation (IPJTD, Methods) in the following, a measure that increases for more jerky trajectories.

**Fig. 2.**
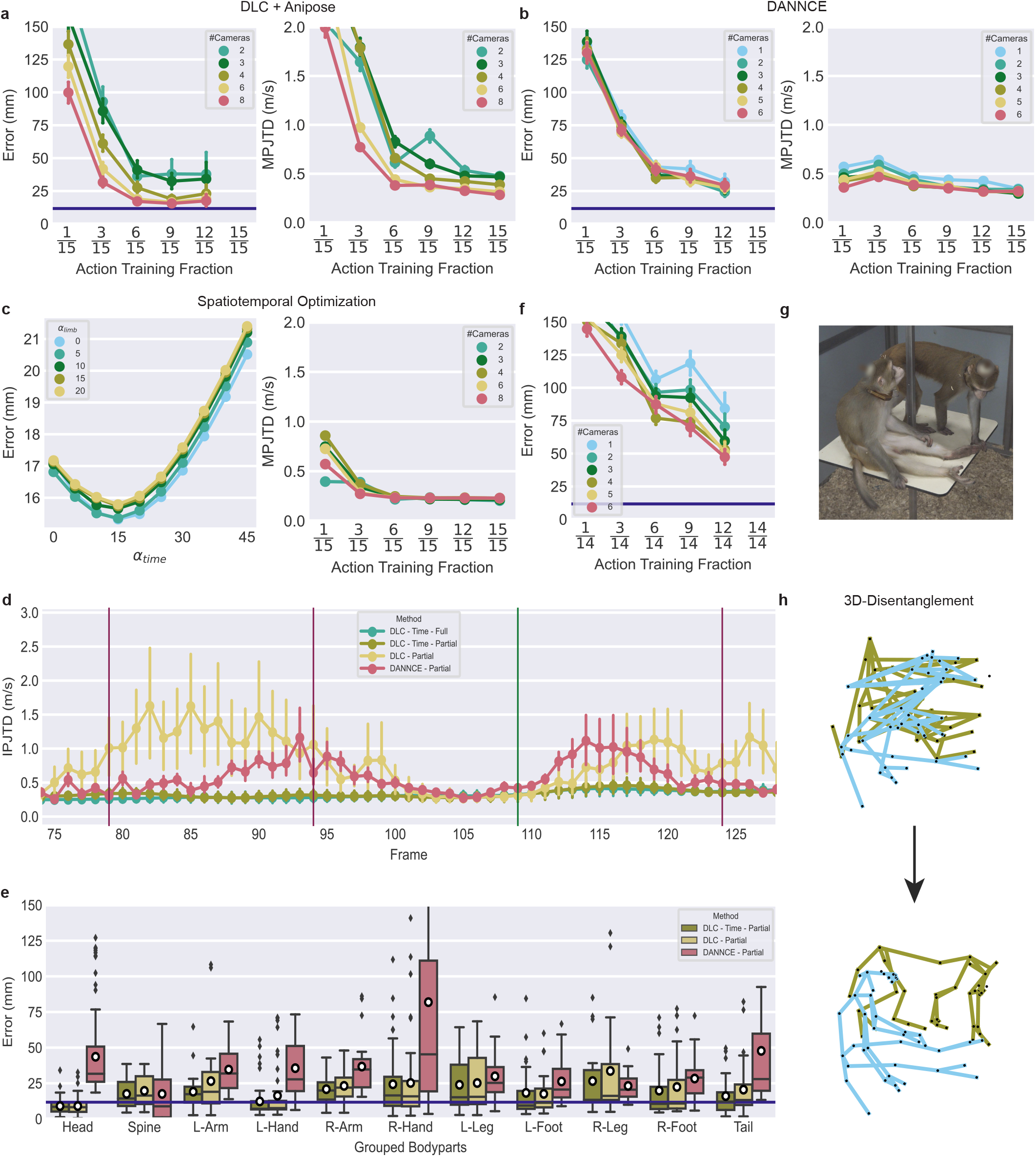
Single-action pose estimation. **a**, Euclidean error on test frames that were not in the training fraction (left) and MPJTD for the entire sequence (right) for DLC with RANSAC and structural refinement. The violet horizontal line indicates the spatial resolution (here, ∼ 11.72mm) of the volumetric approach, DANNCE. In **a-c, d** and **f** points indicate the arithmetic means, and the vertical lines illustrate the 95% confidence intervals. **b**, Euclidean error for equivalent test frames that were not in the training data fraction (left) and MPJTD for the entire sequence (right) for DANNCE for a maximum of six cameras. **c**, Optimization of Anipose’s spatiotemporal parameters for six frames and eight cameras (left). MPJTD for all cameras and training fraction with optimally adjusted parameters for triangulation refinement (right). **d**, IPJTD across recorded frames. The training of the different methods utilized data from six cameras. When only six annotated frames (a subset of all labeled frames of the action) were used, this condition is referred to as ‘Partial’. Conversely, the term ‘Full’ denotes the inclusion of all available labels for these views. The results for DLC are shown with structural refinement only, and applying Anipose’s spatiotemporal optimization. The method setting *α*_limb_ = 0 is referred to as ‘Time’ in **d** and **e**. Labeled frames that were not included in the training set for the partial conditions are indicated by long red vertical lines. The labeled frames that were included in the training set for the partial conditions are indicated by the green vertical line. **e**, Euclidean errors as box plots for keypoints grouped in terms of body parts. The box plot shows the median with interquartile range (IQR), whiskers at 1.5× the IQR and outliers depicted as diamonds. The arithmetic means are shown as white circles. **f**, Euclidean error for multi-animal action prediction with DANNCE. **g**, Image of a labeled frame from the multi-animal condition. **h**, The 3D-Disentanglement procedure removes the confusion of keypoints between the two animals. Triangulated predictions before (top) and after disentanglement (bottom).

We optimized the control parameters of Anipose’s spatiotemporal triangulation refinement by minimizing the error when using six labeled frames and eight cameras (Fig. 2c, left), resulting in significantly smoother trajectories. For optimized parameters, the mean per-joint temporal deviation (MPJTD), a measure for the motion prediction quality [28] given by the average IPJTD over the action, reached a plateau already with only about a third of the number of labeled frames that is established when using DLC and Anipose on all data of the action (Fig. 2c (right),d and Supplementary Videos 3 and 5).

When grouping keypoints by body parts, accuracy was higher for DLC and Anipose with structural refinement than for DANNCE, except for the spine and the mostly occluded right leg, given the same number of cameras and amount of training data. We were able to reduce the remaining errors in DLC by including the optimized temporal constraint (Fig. 2e and Supplementary Fig. 1b). Similar results were obtained for the neutral walking behavior, which was labeled less densely in time (2.87 LPS). For the test frames, DLC and Anipose showed smaller prediction errors compared to DANNCE, both with and without temporal refinement of triangulated landmarks (Supplementary Fig. 1c-e).

We also tracked two monkeys in a sparsely labeled multi-animal action. Since DLC’s multi-animal architecture did not separate markers identities across individuals for so little data, we developed an algorithm (see Supplementary Note) that resolves the assignment of the markers to the individual animals in 3D (Fig. 2g,h). However, this post-processing step was not always robust, especially when DLC’s predictive accuracy of markers was low. We therefore provided DANNCE with 3D input volume locations per individual to implicitly solve the marker prediction and assignment problem. For more labeled keyframes and cameras, 3D keypoint errors were substantially reduced, where including labeled frames from single animal tracking further improved performance (Fig. 2f and Supplementary Fig. 2).

### Single-action reconstruction in Panoptic Studio

For human pose estimation, volumetric landmark detection [27] has been shown to be superior to post-hoc triangulation approaches on the large-scale human dataset Panoptic Studio [29].

To verify if our results transfer from monkeys to humans, we extracted two dynamic actions of a single actor (Fig. 3a) and a group action involving six individuals from this data set. For the single actor, DANNCE, trained with six cameras and labeling half of the frames, reached an average error of 37.9 *±* 33.5mm (Fig. 3b).

**Fig. 3.**
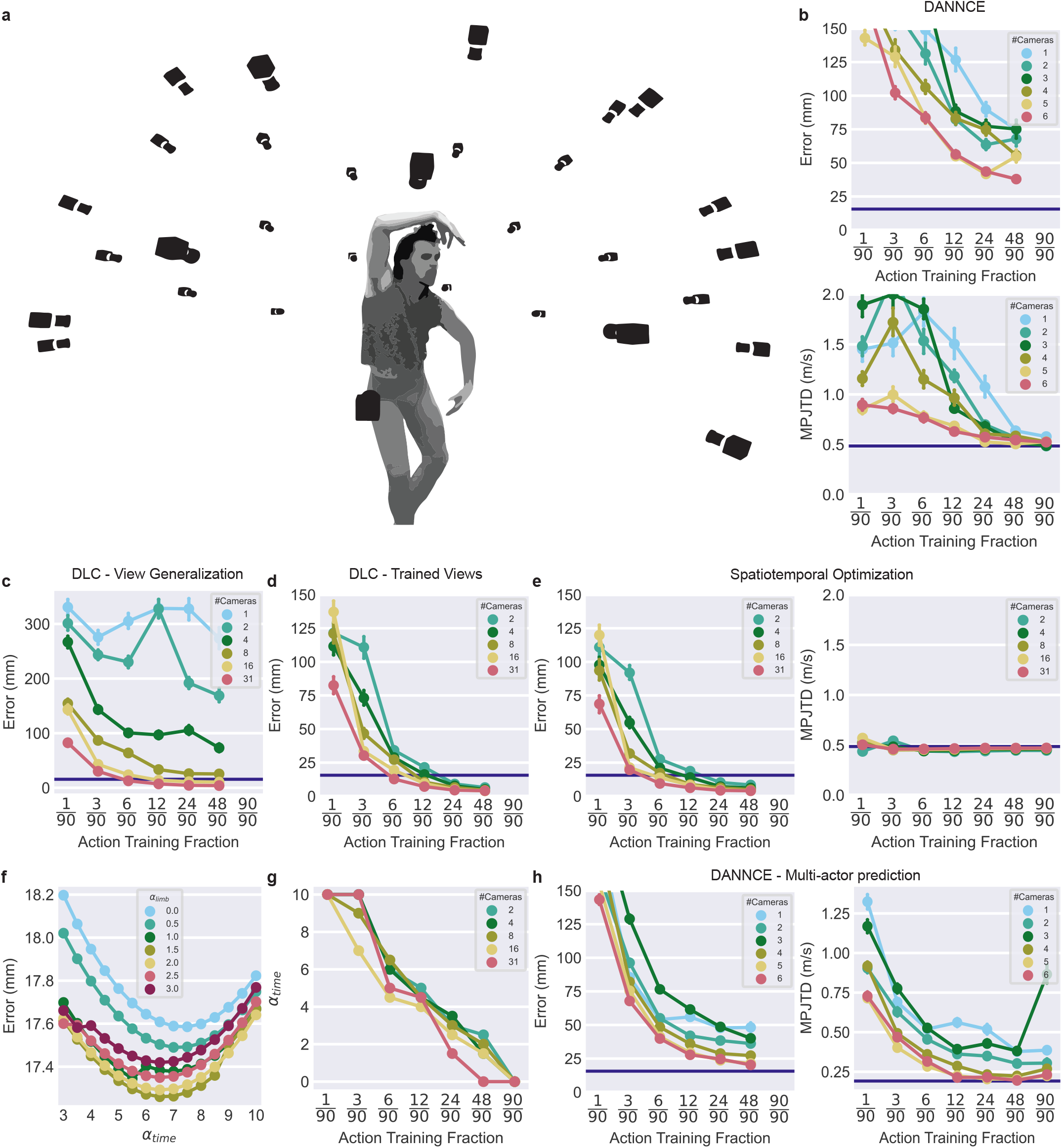
Results for DANNCE and DLC for single-action reconstruction in Panoptic Studio. **a**, Illustration of the Panoptic camera setup: a geodesic dome surrounds actors with 31 HD cameras. **b**, Euclidean error on test frames (top) and MPJTD for the entire sequence (bottom) for DANNCE for single actor action tracking. Horizontal lines in the error plots in **b-e** and **h** indicate the resolution (here, 15.625mm) of DANNCE’s volumetric approach, where horizontal lines in the MPJTD plots in panels **b**,**e** and **h** show the ground-truth mean marker speeds for the single (0.49m/s) and group (0.19m/s) actions. In **b-h** points indicate the arithmetic means, and the vertical lines illustrate the 95% confidence intervals. **c**, Euclidean error for DLC with DLT triangulation when models trained on a subset of camera views were used to predict on all available views. **d**, Euclidean error for DLC with DLT triangulation if models predict on camera views that were present in training only. **e**, Euclidean error (left) and MPJTD (right) for DLC with DLT-based spatiotemporally optimized triangulation. Models predict on camera views that were present in training only. **f**, Optimization of Anipose’s spatiotemporal parameters for six frames and eight cameras. **g**, Optimization of Anipose’s temporal adjustment with an optimized spatial constraint, derived from **f** for different numbers of training frames and cameras. **h**, Euclidean error (left) and MPJTD (right) for the multi-actor action prediction with DANNCE.

**Fig. 4.**
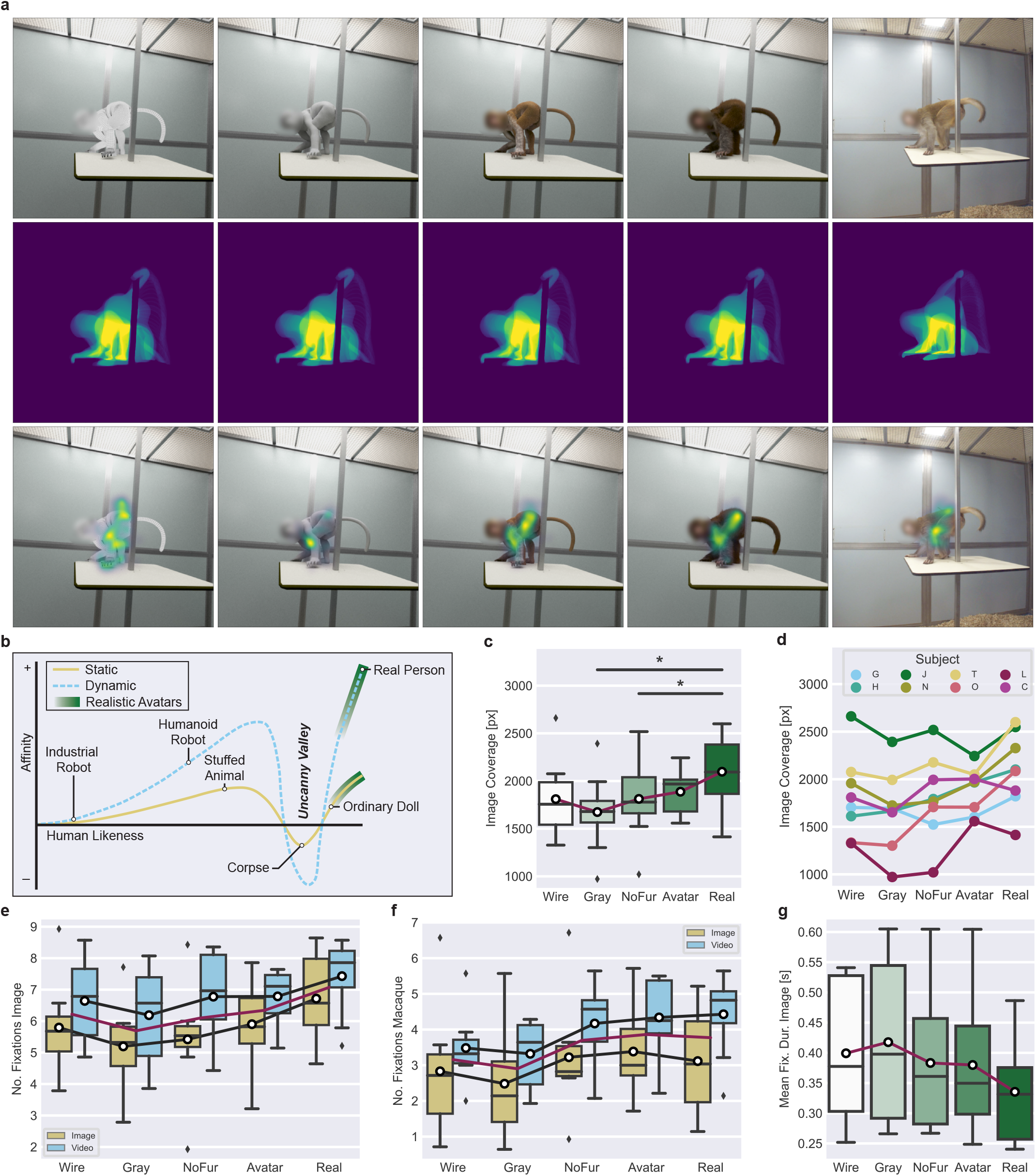
Uncanny valley effect for bodies in macaques. **a**, First row: stimuli for the uncanny valley experiment with different degrees of realism, increasing from left to right. The rightmost stimulus is a real video. Second row: superposition of the monkey silhouettes (for degraded stimuli derived from the avatar with fur) across all frames. Last row: accumulated fixations falling into these dynamic masks for the eight monkey subjects across all trials. **b**, Illustration of the uncanny valley hypothesis in human perception, adopted from [18]. The range of realistic avatars, both static and dynamic, is shaded in dark green. **c**, Box plot of image coverage by fixations in pixels for different render types. Plots in **c-f** show the results of the first trials only, while panel **g** shows the analysis including all trials. The box plots show the median with interquartile range (IQR), whiskers at 1.5× the IQR, and outliers depicted as diamonds. The arithmetic mean is indicated by a white circle. In **c** and **e-g**, arithmetic means are connected by a red line. Statistical significant pair-wise differences are indicated by horizontal lines accompanied with asterisks: *p*^*^ *<* 0.05 and *p*^**^ *<* 0.01. **d**, The mean coverage shown as a function of render type and for the eight monkey subjects shown individually. **e**, The number of fixations on the image as a function of render type, shown separately for two presentation types: static (image) and dynamic (video). In **e** and **f**, arithmetic means per condition are connected by black lines, with the overall mean per render type indicated by a red line, omitting the additional white circles. **f**, The number of fixations within the macaque silhouette for the different render and presentation types. **g**, Box plot of the mean image fixation duration, for each render type and including all trials.

In comparison, DLC achieved a similar error level even with only four cameras and six labeled frames (28.8 *±* 47.6mm), where we utilized the direct linear transformation (DLT) for triangulation (Fig. 3b, d). Including additional camera views in the prediction that were not part of the training data increased the error rather than reducing it (Fig. 3c). After optimizing the spatiotemporal parameters of Anipose’s triangulation refinement, the prediction error fell below the resolution limit of DANNCE’s input volume of 15.625mm (14.0 *±* 17.1mm; Fig. 3e,f), even for a small number of cameras and labeled keyframes.

DANNCE matched the mean marker speeds of ground-truth labels only with substantial amounts of training data (Fig. 3b). In contrast, DLC required a lower amount of training data to reach a comparable level of trajectory smoothness (Supplementary Fig. 3a). When fixing the spatial constraint in triangulation, the temporal adjustment could be decreased as more frames were included in the training set (Fig. 3g).

For tracking multiple individuals, the errors for DANNCE were lower than for tracking single individuals (20.3 *±* 20.2mm). This was accompanied by matching the ground-truth mean marker speed with less training data, likely since labeling six actors results in six times as many labels per keyframe (Fig. 3h).

Adding frames from the single-actor action to the training dataset did not yield improvements (Supplementary Fig. 3b). Consistent with our findings for macaques, tracking a single action with limited data was more accurate using the triangulation approach, while an inclusion of more cameras and keyframes significantly reduced the tracking error for multiple agents for DANNCE.

### Uncanny Valley for body perception

To investigate the extent to which the animated avatar is perceived as realistic by real animals, we recorded eye movements of eight macaques, while they observed images and videos of an acting monkey or monkey avatar. We generated stimuli with intermediate degrees of realism (Fig. 4a) for this uncanny valley experiment from an emotionally neutral walking and a submissive body action, building upon previous work focused on faces [5].

The animals’ behavior was quantified by eye-fixation analysis. We defined areas of interest (AOIs [30]) based on the monkey silhouette that was tracked using Track-Anything [31]. We then quantified visual exploration by the stimulus coverage in pixels, derived from eye fixation positions. A group-level analysis of the first trials revealed an uncanny valley in visual exploration (one-way repeated measure analyses of variance (ANOVA): *F* (4, 28) = 6.953, *p* = 0.002, Fig. 4c,d and Supplementary Table 1), with the furless and particularly the gray avatar being fixated least frequently, resulting in the U-shaped dependence of looking preference as a function of the degree of realism of the stimuli.

Across all trials, the uncanny valley shape was less pronounced for the number of fixations on the entire image, the macaque silhouette, and especially for image coverage (Supplementary Fig. 4a,d). These changes in gazing behavior were accompanied by a decrease of total fixation counts with increasing number of trials (Supplementary Fig. 4f), potentially reflecting a decay in the animals’ attention to the stimuli. However, the average fixation duration across all trials and subjects showed a reversed uncanny valley where animals looked longer at less realistic stimuli (Friedman: *χ*^2^(4, *n* = 8) = 19.3, *p* = 0.0007, Fig. 4g). The same trend was observed for fixations falling into the monkey silhouette (Friedman: *χ*^2^(4, *n* = 8) = 13.8, *p* = 0.008, Supplementary Fig.4b). Analyzing only the fixation duration of the first trials, the reversed uncanny valley was less prominent for whole image and monkey shape fixations (Friedman: *χ*^2^(4, *n* = 8) = 11.8, *p* = 0.019, Friedman: *χ*^2^(4, *n* = 8) = 6.9, *p* = 0.14, Supplementary Fig.4c). Overall, these findings demonstrate that monkeys exhibit an uncanny valley effect for bodies. While such effects have been reported for facial stimuli [5, 20], this is the first demonstration of this phenomenon for bodies in monkeys.

### Influences of motion and action

The influences of the different factors in our experimental design were further quantified by generalized linear models (GLM), assuming negative binomial distributions (Supplementary Table 2) to model the response variable, that is, counts of whole image and monkey shape fixations. For both, likelihood-ratio tests for data from initial trials show a significant influence (*p <* 0.0001) of the factors Render and Presentation type (dynamic vs. static). Yet, the interaction of these two factors is not significant (*p* = 0.5502 and *p* = 0.8086, Supplementary Tables 3,4). Except for the uncanny valley of macaque silhouette fixations in static images (Friedman: *χ*^2^(4, *n* = 8) = 7.465, *p* = 0.1133 (images), *χ*^2^(4, *n* = 8) = 9.4842, *p* = 0.0050 (videos), black curves in Fig. 4f), we found a significant influence of the factor Realism, also for both images and videos in whole image fixations (Friedman: *χ*^2^(4, *n* = 8) = 10.843, *p* = 0.0284 (images), *χ*^2^(4, *n* = 8) = 12.662, *p* = 0.0131 (videos), Fig. 4e). Subsequent post-hoc analyses, however, did not indicate significant differences between the different levels of Realism for either Presentation type.

Consistent with the classical uncanny valley hypothesis (Fig. 4b), where dynamic presentations are preferred except deep within the uncanny valley, videos were overall fixated more often than static images.

Furthermore, the macaques’ gazing behavior was consistent across actions when presented with different degrees of realism. In particular, the neutral action accounted for more fixations than the submissive action in both images and videos (Supplementary Fig.4e, Supplementary Table 5).

We also confined the analysis to real footage and the most realistic avatar. In this case, the GLM analysis revealed a significant effect of the displayed action, while the factor Render (real macaque vs. avatar) was not significant, nor were the interactions (Supplementary Table 6). Thus, macaques responded very similarly to the avatar and the real video, showing that the developed method creates animations that appear to be realistic to macaque monkeys.

## Discussion

Here we presented MacAction, a tracking and animation system that combines accurate 3D markerless motion capture with the animation of a highly realistic macaque avatar. Animating such complex computer-graphics models necessitates tracking numerous markers in 3D. In contrast to human tracking, for which extensive databases with precise kinematic data are available, the creation of similar databases for monkeys is not economically feasible.

To solve this problem, we established a pipeline for video-based tracking with high accuracy, requiring minimal hand-labeled data for individual actions. We demonstrated that labeling two frames per second is sufficient to generate realistic 3D animations for complex body actions. Especially for larger marker sets, as required in computer animation, our method significantly reduces hand-labeling compared to established methods in 3D behavioral analysis [9, 10, 26, 32].

At the same time, our approach tracked single actions more precisely than other behavioral tracking approaches in macaques, even ones including additional sensor devices such as RGB-D cameras [21, 33].

For the tracking of individual animals with little data, we further obtained better results with a method using post-processed triangulation of 2D marker trajectories than for 3D volumetric pose estimation. For the simultaneous tracking of multiple animals, the volumetric approach was more practical, and further improvement can be expected from incorporating temporal supervision [28]. Considering the known challenges of tracking macaques [7, 9], our method is likely adaptable to other animals with simpler kinematic structures. This is particularly needed for studying actions of endangered animal species [34], where data might be scarce and costly to obtain.

MacAction goes significantly beyond a new approach for motion tracking, since it includes the entire process of crafting digital avatars, motion retargeting, scene asset creation, rendering, and ultimately, validating animations with real monkeys. Our eye-tracking data from behavioral experiments showed that macaques respond to our dynamic avatars similarly to actual video recordings, proving the high degree of realism in the generated animations.

By exploiting eye movement analysis methods that have been involved in demonstrating an uncanny valley in monkey face perception [5, 20], we establish the first uncanny valley effect for the observation of bodies in macaques. The fact that humans also show uncanny valley effects for bodies [35, 36] supports the hypothesis that neural mechanisms for processing visual stimuli of bodies are shared among different primate species. The future study of macaque monkeys with such highly controlled and realistic visual stimuli will be crucial for unraveling the underlying neural principles of social vision.

## Methods

### Animals and husbandry

Eight male rhesus monkeys (Macaca mulatta), ranging in age from 5 to 7 years, contributed to this study. The animals are housed in enclosures at the KU Leuven Medical School and experience a natural day-night cycle. Each monkey shares its enclosure with at least one other cage companion. On weekdays, dry food is provided ad libitum, and the monkeys obtain water, or other fluids, during experiments until they are satiated. During weekends, the animals receive water along with a mixture of fruits and vegetables. The animals have continuous access to toys and other forms of enrichment. For experiments unrelated to this paper, the monkeys were implanted with a plastic headpost, attached using ceramic screws and dental cement following standard aseptic procedures and under full anesthesia. Both the animal care and experimental procedures adhere to regional (Flanders) and European guidelines and have been approved by the Animal Ethical Committee of KU Leuven.

### Multi-camera recording setup

We created a 3D model of the surrounding scene in the animal house using a commercial imaging laser scanner (LEICA BLK360). The resulting extracted colored point clouds from multiple scans were registered using Scasa PinPoint (v2.6.0), and further processed with Autodesk Recap 2022. The size of the surrounding cage was (2.1 m x 3.1 m x 2.3 m). Based on these scans, we reconstructed the entire surrounding environment as a 3D model in Maya 2022. The reconstructed scene elements were then imported into the real-time gaming platform Unreal Engine (4.27.1). In this software environment, we created virtual cameras to emulate those we planned to use in the final recordings. Adjustments to these cameras, including their extrinsic position and field of view (FOV), helped to choose appropriate lenses. We determined the focal length in relation to the film size in order to achieve a suitable FOV. We then verified that for the chosen lenses a virtual cube with a side length of 75 cm, which represented the macaque’s movement space, was fully captured in the FOV. This cube was centrally placed in the virtual cage, and it was captured at a resolution of 2056x2056 pixels by the virtual cameras.

For the real recordings, we employed eight high-resolution machine vision color cameras from IO Industries. Four cameras of the type Victorem 51B163CCX (2464x2056 pixels) and four of the type Victorem 120B68CCX (4112x3008 pixels) were initially positioned at the virtual camera locations. For synchronization and recording, we used two digital video recorders (CORE2CXPLUS) from the same manufacturer, enabling precise synchronization with low-latency TTL signals. Together with the cameras also eight LED light panels (StroboMini2, Norka Automation) were controlled by TTL signals. This allowed to lit the scene in synchrony with global camera shutters, reducing the overall brightness experienced by the monkeys significantly. As a result, the cameras’ exposure time could be set to as low as 300 microseconds, keeping motion blur effects at a minimal level. Four light panels surrounded the cage horizontally, and the remaining ones were placed on top of the cage next to a single camera, recording a top view. The cage’s top consisted of a metal grid on which a plexiglass plate was mounted, protecting the cameras from manipulations by the animals. The remaining equipment was mounted on tripods outside of the cage. Since the setup was not completely protected against mechanical perturbations, we re-calibrated the optical recording system regularly before each recording session using two different calibration schemes. The first scheme involved a custom ChArUco pattern (39.7cm x 39.7cm, 0.5 marker-to-square ratio, OpenCV standard 4-bit dictionary with 50 markers for ArUco checkers). This pattern was detected by software written in Python (3.9.7) using OpenCV (opencv-python 4.5.5.62 and opencv-contrib-python 4.5.5.62). Additionally, the extrinsics were optimized with bundle adjustment in MATLAB (R2021b). The second calibration method involved a light-emitting wand from Vicon (Active Wand), developed for the calibration of a commercial marker-based motion capture system, and part of proprietary post-processing software. Both methods achieved subpixel accuracy. After the recordings, we transferred raw video material at high speeds to an external storage system (D5 Thunderbolt 3, Terramaster) with up to 90TB storage capacity.

### MacAction Software

MacAction was developed using Python 3.9.7 and incorporates several widely used open-source software packages for the scientific community: numpy 1.22.1, aniposelib 0.4.3, matplotlib 3.5.0, pyqtgraph 0.11.0, opencv-contrib-python 4.5.5.62, opencv-python 4.5.5.62 and scipy 1.7.3. We employed established keypoint estimation software, anipose 1.0.1, deeplabcut 2.2.2 (cuda 11.4 and cudnn 8.2.1) and dannce 1.2.0 (cuda 10.1 and cudnn 7.6.5). Our repository includes code modules for various functions: video compression, gamma correction, camera calibration, and calibration file conversion. We further developed and provide the package AnimLabel, a labeling tool to generate 3D poses from simultaneously presented multi-view images. AnimLabel enforces consistent labeling across an arbitrary number of views by including multi-view geometric constraints from calibration data. Epipolar lines are utilized to reduce relevant regions across views to a single line or intersections of lines if a keypoint is labeled in a single or multiple views. To further reduce the labeling effort, we project triangulated landmarks of at least two views into all other perspectives. In practice, we use at least three hand-labeled markers to reduce the uncertainty of the triangulated 3d keypoint positions. In this way, labeling effort remains the same, but labeling throughput scales linearly with the number of cameras. We also ensure sufficient labeling accuracy by setting a threshold for the mean reprojection error across labeled views (5 pixels).

The software provides the possibility to crop views, showing only relevant image areas, which is especially relevant for high-resolution images. AnimLabel can also export cropped single animal actions per view. Based on a pre-defined spatial margin, we project all markers into the image plane. We then identify the markers that are most distant from the center of the frame to costruct a cropped frame that does not clip any of the body regions.

Data labeled through AnimLabel, can be exported in appropriate formats to both, DANNCE and DeepLabCut for multi-view and multi-animal projects. We convert the tracking data from DeepLabCut to Anipose’s format for single- and multi-animal tracking. All generated full-body trajectories are then saved in a numpy binary format (’.npy’). For rendering, we use Maya, leveraging its native Python 3 interpreter, mayapy, which allows to import and retarget motion data. Maya allows to enhance the quality of the motion capture data by interpolation to smoother trajectories, and we exploited calibration data to align virtual cameras with their real-world counterparts.

### Datasets

Our study generated or used the following datasets:

#### Macaque animation

Video data of macaques was recorded at 80 frames per second (FPS). From 2.5 hours of recording data, we extracted three distinct actions: a submissive action (3.43s), a neutral action (2.44s), and an interaction of two macaques (93.4s) with 15, 7 and 14 labeled frames, respectively. Based on these labeled frames, we assessed the accuracy of our tracking pipeline in three different scenarios: (1) oversampling a short, complex body movement; (2) providing fewer labels for a concise, simpler action; and (3) fewer labeled time points of a prolonged multi-animal close-interaction sequence.

We labeled an extensive set of 42 body landmarks: ‘Jaw’, ‘lowerLip’, ‘upperLip’, ‘lCornerM’, ‘rCornerM’, ‘Nose’, ‘lEye’, ‘rEye’, ‘Head’, ‘lEar’, ‘rEar’, ‘Neck’, ‘lShoulder’, ‘lElbowOut’, ‘lElbowIn’, ‘lWristOut’, ‘lWristIn’, ‘lThumb’, ‘lMidK’, ‘lMidTip’, ‘rShoulder’, ‘rElbowOut’, ‘rElbowIn’, ‘rWristOut’, ‘rWristIn’, ‘rThumb’, ‘rMidK’, ‘rMidTip’, ‘mid-Spine’, ‘lHip’, ‘lKnee’, ‘lAnkle’, ‘lBigToe’, ‘lFootTip’, ‘rHip’, ‘rKnee’, ‘rAnkle’, ‘rBigToe’, ‘rFootTip’, ‘beginTail’, ‘midTail’, and ‘tipTail’. This allowed us to infer both the positions and rotations of individual joints in 3D, crucial for controlling a high-quality virtual avatar.

Due to the significant amount of work involved in labeling markers on larger data sets, we limited the number of keypoints to the minimum necessary for animating the chosen avatar model.

#### Human Benchmark

To evaluate the amount of labeling for multi-view tracking approaches used in this paper, we also utilized a publicly available large-scale data set for humans, the CMU Panoptic dataset [29]. Due to the large number of cameras provided by the dataset, it has significantly contributed to human pose estimation and has led to the development of volumetric triangulation [27], which forms the basis of the DANNCE method [24].

We refrained from evaluating our method on datasets constructed with physically attached markers ([37], [38]). These keypoints can become salient in images and skew the learning process for markerless tracking. In contrast, the Panoptic Studio dataset provides motion capture data without physical markers, utilizing recordings from 480 VGA, 31 HD, and 10 Kinect v2 RGB+D cameras, evenly distributed in a geodesic dome. Designed to capture human social interactions, it comprises actors engaged in activities such as playing games and instruments, or showing various other types of movements. For comparison with our monkey dataset, we extracted short sequences from movies showing two actions: a complex dance movement, and a clip involving six individuals engaged in simultaneous interactions. Each action consists of 90 frames lasting approximately 3s. We used data from up to 31 HD cameras at a resolution of 1920x1080 pixels.

The volumetric approach of the DANNCE algorithm has the disadvantage of a limited tracking volume. In order to better align the algorithm’s accuracy between humans and macaques, we restricted the included 3D poses to upper body keypoints, and hips. Consequently, the human marker set comprised the following 15 keypoints: ‘Neck’, ‘Nose’, ‘BodyCenter’, ‘lShoulder’, ‘lElbow’, ‘lWrist’, ‘lHip’, ‘rShoulder’, ‘rElbow’, ‘rWrist’, ‘rHip’, ‘lEye’, ‘lEar’, ‘rEye’, and ‘rEar’. Finally, we sampled actions over time and across different numbers of cameras to create diverse datasets for benchmarking the tracking algorithms.

### Pose estimation

*Single animals*. For single-animal pose estimation with DeepLabCut 2.2.2, we fine-tuned a pre-trained ResNet-101 model on the ImageNet dataset using default training configurations. We created the dataset by exporting actions from AnimLabel. Specifically, we included both single and multiple animal actions across views in a label file matching DLC’s structure, bypassing its graphical user interface using custom scripts.

After training and inference on a particular view, we consider all 2D keypoint predictions, regardless of their confidence values. In this way, we do not restrict information prior to the triangulation process. We use Anipose’s RANSAC triangulation method and optimize the 3D points by spatiotemporally regularized triangulation, an extension that incorporates not only the reprojection error [26]:

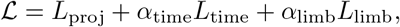

where the respective loss *L*_proj_ aims at minimizing the distance between the projected 3D markers and the associated 2D predictions over cameras. Since 3D body marker trajectories should be smooth in time and ensure consistent limb lengths, the losses *L*_time_ and *L*_limb_ are added, which measure the mean joint deviations over time and with respect to the expected limb lengths between connected keypoints. The parameters *α*_*i*_ in the composite loss allow to adjust the importance of these different factors in the cost function. We refer to the triangulation algorithm for the choice *α*_time_ = *α*_limb_ = 0 as structural refinement, as initial 3D points are optimized and shifted only to yield a consistent projection across views.

For DANNCE, we fine-tuned pre-trained weights of their AVG network architecture on a six-camera setup for rat pose estimation. As with DeepLabCut, we created custom scripts to align the labeling structure with AnimLabel and used the default training configurations. The algorithm requires a prediction of the animal’s 3D center of mass (COM), which was determined using quadratic B-spline interpolated hand-labeling (of the ‘midSpine’ for macaques, and ‘Neck’ for the upper human body), instead of training a separate COM network within DANNCE. We specified the encapsulating volume size by determining the volume that encompassed all individuals and poses in the dataset, adding an additional safety margin. We did not incorporate any temporal or spatial smoothing, except for potential interpolation of keypoints in the commercial animation software.

#### Multiple animals

For testing the multi-animal tracking using DLC, we incorporated an additional re-identification step. This step associates the estimated 2D keypoint trajectories with the corresponding individual, exploiting multi-view consistency constraints. Furthermore, we introduced spatial and temporal smoothness constraints to derive a continuous re-identification of individuals (see Supplementary Note). The re-associated 2D projections are then transferred to Anipose’s optimization pipeline, where we treat multiple individuals as a single entity without connecting respective skeleton architectures.

With DANNCE, we did not change the overall network architecture as with DLC, but only the COM location in 3D to probe the network. Specifically, each of the two animals is associated with a different COM location. As with single animals, we automatically adapted labeling structures for multi-animal tracking.

#### Error metrics

To quantify the position accuracy of the tracking, we use the Euclidean distances between the predicted and the ground-truth 3D joint positions. As in [27, 28], we computed the mean per-joint position error (MPJPE):

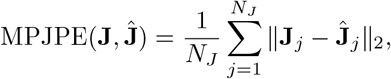

where the vector **J** includes the *N*_*J*_ ground-truth 3D keypoint coordinates, and **Ĵ** denotes the corresponding tracking estimate. We average this per-frame error across the entire action to provide an overall error measure for the sequence. In addition, we compute Euclidean errors for specific body parts in the same way, including only the landmarks of those body parts.

Another measure characterizes the smoothness of the tracked trajectories. If the tracking fails, typically the trajectories show jumps and non-smooth transitions, for example, when the triangulation procedure misaligns 2D keypoints across cameras. Inspired by [28, 39], we define a measure for the temporal smoothness of the predicted action of length *T* by the mean per-joint temporal deviation (MPJTD):

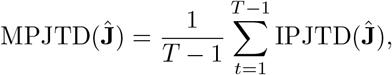

the action’s average of the instantaneous per-joint temporal deviation (IPJTD):

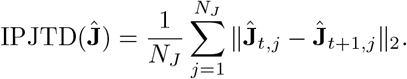

The IPJTD can be thought of as the instantaneous speed or first-order derivative of the predicted keypoints, where large amounts and jumps are more likely due to erroneous tracking than fast movement.

### Animation

*Model*. We utilized a commercial macaque avatar model from CGTrader (https://www.cgtrader.com/3d-models/animals/mammal/monkey-fur-rigged). This digital macaque model consists of a mesh with 54,806 vertices, includes 8192x8192 pixel resolution textures, and fur. It is fully rigged, i.e., linked to a hierarchical structure of 86 joints that allows for posing the character with appropriate deformations of the surface structure. It uses linear blend skinning to determine these surface deformations. To enhance the realism of the avatar, we scaled hands and feet to match measurements of one real monkey, derived from triangulated poses of hand-labels. Preliminary eye-tracking results indicated that the animals were specifically interested in the rear part of the observed bodies. Consequently, we refined this area of the model by attaching a pair of testes and two patches of keratinized skin (sciatic protuberances), characteristic of Old World monkeys, using rivet constraints in Maya 2023. Additionally, we utilized the XGen Interactive Groom Editor to modify the fur in this area, aiming for a less dense and shorter fur texture.

#### Pipeline

Utilizing the tracked 3D keypoint trajectories of all markers, we implemented a customized yet generally applicable interface to the kinematic tree of a digital avatar model in Maya. The keypoints are rescaled to match the dimensions of the avatar, and the character’s joint positions and rotations are set to their default states, known as the rest pose. We then position the root joint of the avatar in the scene, and determine the rotations of individual joints along its kinematic tree by solving Euler angle transformations in matrix form consecutively. A detailed description of the approach can be found in the Supplementary Note. In this process, we distinguish between joints of the avatar model with two and three rotational DOFs. For body segments in the avatar with more intermediate joints than tracked keypoints, that is, the spine and tail, we perform a quadratic B-spline interpolation, treating these joints as having two DOFs.

### Benchmarking details

All benchmarking and training procedures were conducted on Nvidia Geforce RTX 3090 and 4090 GPUs. In the datasets, we prioritized cameras that increase view variability.

#### DLC

We trained a single model across views for each species. Single-animal models were trained for 5000 epochs and multi-animal models for 1000 epochs. All models utilized a ResNet-101 backbone. Data sets with more than five images were divided into 95% training, and 5% testing sets. After training, we predicted 2D landmarks on complete video sequences. These predictions were then converted into 3D trajectories using Anipose.

#### DANNCE

Our training followed the guidelines provided on https://github.com/spoonsso/dannce/wiki. We trained models across different animal types and dataset sizes, each for 1000 epochs. Due to limited data sizes, we excluded validation samples. For all species, models were trained using a 64x64x64 volume grid, resulting in voxel resolutions of 11.719 mm (macaques) and 15.625 mm (humans), based on the cubic volume side lengths of 750 mm and 1000 mm, respectively. For training configurations with less than six cameras, models trained on duplicated camera views.

### Uncanny Valley Stimuli

For the behavioral experiment testing the quality of our animated macaque avatar, we adopted a similar approach as in experiments that showed the presence of an uncanny valley in macaques for faces [5]. According to the uncanny valley hypothesis, variations in the avatar’s realism should induce changes in preference, which can be discovered by eye tracking. Starting from the complete avatar model, we created three degraded versions of it: (1) discarding the fur of the model and depicting the avatar with skin texture only; (2) removing the color information of the skin by the replacement of a smooth gray texture; and (3) discarding light reflection properties of the surface structure by outlining the mesh’s edge lines in black on a constant white background. As actions for these stimuli, we used the emotionally neutral walking and the submissive body movement.

For the final animations, we also reconstructed the background scenery, including the cages and lights, as 3D scan-based models of the individual geometric elements in Maya. To reproduce the recording conditions virtually, we positioned rendering cameras according to their real-world counterparts. Due to the remaining shape difference between the real macaque and the avatar, the same motion can result in slightly different trajectories of the corresponding body parts, most apparent in the distal parts of the avatar’s limbs. To reduce these differences, we adjusted strongly deviating motion paths in Maya 2023.1 using the Graph Editor tool, ensuring a realistic physical interaction of the avatar with its environment.

Excluding the unnatural viewpoint from the top of the cage, we rendered seven different viewpoints per action and avatar configuration using V-Ray for Maya (v5.20.02). In line with rendered videos, we adjusted real videos to square format and dropped every third frame. As a result, the uncanny valley dataset comprises 70 dynamic stimuli derived from two actions, seven viewpoints, and five levels of realism (including a real video). To distinguish influences of motion and shape in the uncanny valley experiment, we selected from each action a key pose that best represents the particular expression, and extracted the corresponding still images from the videos. This yielded a final set of 140 stimuli, half static and half dynamic. Given the significant influence of the face on eye fixations [40, 41], visible faces were blurred using a Fast Box Blur filter in Adobe After Effects 23.1.0 (https://www.adobe.com/de/products/aftereffects.html). Additionally, parts of the real video showing researchers or body reflections were blurred, ensuring equivalent blurred areas in the rendered videos. The mean luminance of all videos and images were equated, taking into account the gamma function of the display using custom written software.

### Eye tracking

#### Setup and paradigm

During the experiment, subjects were seated in a primate chair with their heads fixed. For eye tracking, we used an infrared eye tracker (1000 Hz EyeLink CL, SR Research) sampled at 2000 Hz in front of 22.5-inch, high-resolution LCD screen (VIEWPixx, 1920x1200 pixels, 120 Hz refresh rate), specifically designed for eye fixation experiments. To map the monkey’s gaze to the image’s pixel space, we conducted a calibration experiment at the beginning of each recording session. In this experiment, a red square fixation target (size = 0.2°) was presented on a gray background at 13 different locations, specified by the x, y coordinates in percentages of the display extent: [10,5], [50,5], [90,5], [35,30], [65,30], [10,50], [50,50], [90,50], [35,70], [65,70], [10,95], [50,95], [90,95].

In each trial, the target appeared at a new location, and the monkey received a juice reward for making a saccade towards the target and maintaining fixation for either 800 (for subjects: C, J, L, N and O) or 1000 ms (for monkey G, H and T). The duration of fixation varied depending on the subject’s level of training. On average, we collected 11.9 valid trials at each location, with a minimum of 6 trials. The medians of these fixations at each location were then used to transform gaze coordinates into pixel coordinates, using second-order polynomial transformation (fitgeotrans, MATLAB R2020b). In the subsequent free-viewing test all stimuli had a size of 13.5x13.5° and were presented in a random order for 2.4s with an inter-stimulus interval of 400 ms. The stimuli were centered on the screen and displayed on a gray background having the same luminance level as used during calibration.

There was no fixation target, and the monkeys received a juice reward as long as their gaze remained on the screen, regardless of where they were looking. During a block, each stimulus was presented once. In case the monkey’s gaze left the screen area, the stimulus remained on the screen and was not repeated during the same block to minimize the influence of familiarity on the viewing behavior. At least three runs were included from each subject in the analyses, all recorded on the same day. The start and end of the stimulus presentation were signaled by a photodiode detecting luminance changes of a small square located in the corner of the display (invisible to the animal) and placed within the same frame as the stimulus events. Stimulus presentation, event timing, and juice delivery were controlled by an in-house-developed digital signal processing-based computer system, which also sampled the photodiode signal besides vertical and horizontal eye analog signals.

#### Analysis

TrackAnything [31] (https://github.com/gaomingqi/Track-Anything) was used to define and track appropriate regions for the eye tracking analysis. This software ran locally on a machine equipped with an NVIDIA GeForce RTX 3090. We tracked masks for all real and avatar stimuli videos. Avatar masks were also used for its degraded variants, since fur-less avatars covered the same or less area in the image. For static images, we extracted the relevant masks out of the segmented videos. Eye tracking data from individual subjects were then visually inspected in MATLAB R2021b to define proper values for fixation analysis with the toolbox EyeMMV [42]. The spatial constraint parameters for the clustering of the fixations were both set to ∼ 0.46^*°*^ in visual degrees.

We required a minimum fixation duration of 180 samples or 90 ms, in line with literature on monkey eye tracking, where fixation duration thresholds varied from 50-150 ms [5, 43–46]. Inspired by commonly used human eye fixation metrics [30], we defined three measures for a quantification of differences between the conditions: (1) mean fixation duration; (2) number of fixations that fall into dynamic or static body masks, where in videos the fixation was required to be within body masks for the whole fixation duration to be accepted; (3) coverage or stimulus exploration, *C*. For computing the coverage, we aggregated isotropic Gaussian kernels (*σ* = 9 pixels ≙ 0.244^*°*^) located at eye fixation locations that lay within the image or body shape, represented by the following function over pixel coordinates (*x, y*):

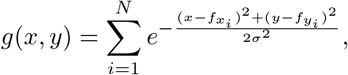

where *N* denotes the total number of fixations for a stimulus, and 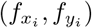 is the location of an individual fixation. We then counted image locations, where this function exceeded a fixed threshold:

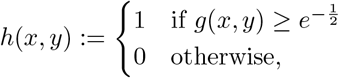

resulting in the coverage:

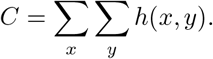

For displaying fixations on stimuli (Fig. 4a), we added up the Gaussian kernels at the eye fixation locations, weighted by their respective fixation durations.

### Statistical analysis

*Tracking*. Euclidean error plots show mean marker Euclidean distances and .95 confidence intervals as vertical lines. Outliers in boxplots are shown as diamonds, and mean values as black outlined white circles. When referring to a specific training configuration in the text, which is characterized by the algorithm, cameras, and labeled keyframes used for training, we present the Euclidean errors in terms of their mean and standard deviation.

#### Behavioral experiment

We performed one-way repeated measures ANOVA after checking residuals of a linear model for normality and outlier tests in R 4.3.1. Afterwards, we used the rm anova() function from pingouin 0.5.3 in Python 3.9.7. If Mauchly’s test of sphericity was significant, we applied a Greenhouse-Geisser correction. Effect sizes are reported as generalized 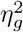. To identify pairwise differences, we adjusted post-hoc pairwise, two-sided t-tests with Benjamini-Hochberg false discovery rate correction.

To model fixation counts within the whole image and the monkey sihouette, we additionally constructed generalized linear models with Poisson and negative binomial distributions using the package MASS 7.3.60 in R. In line with [47], we also constructed generalized linear mixed effect models (GLMMs) with Poisson and negative binomial distributions with the package lme4 1.1.33. Subject differences in fixation counts were accounted for by a random intercept for each individual.

We employed negative binomial GLMs as these showed no effects of under- and overdispersion, unlike all other models. Therefore, we tested fitted model residuals and simulated residuals with the function testDispersion() and inspected Q-Q plots to assess model fit, using the package DHARMa 0.4.6. Afterwards, we computed likelihood-ratio tests (LRTs) with the package lmtest 0.9.40 for different nested parameter combinations to identify interactions within predictor variables.

We used the non-parametric Friedman test from the scipy.stats 1.9.0 package when normality assumptions were not satisfied, or as an additional test for fixation counts. In cases where a significant effect was detected, we performed post-hoc analyses utilizing the Dunn’s test from the package scikit posthocs 0.7.0, applying a Benjamini-Hochberg false discovery rate correction.

To identify pairwise differences in fixation counts when an action or presentation type variable is varied, we utilized two-sided continuity corrected Wilcoxon signed-rank tests from the package pingouin 0.5.3. We report effect sizes in terms of the common language effect size (CLES).

## Data availability

Motion capture data, renderings, eye movement recordings and the virtual scene will be made publicly available upon publication.

## Code availability

The MacAction software package will be made publicly available upon publication.

## Acknowledgements

Funding was provided by the European Research Council (2019-SyG-RELEVANCE-856495). The authors thank the International Max Planck Research School for Intelligent Systems (IMPRS-IS) for supporting Lucas M. Martini. We are grateful to S. Polikovsky for helping in the motion capture hardware design and to D. Cicchetti for support in labeling actions. We thank I. Puttemans, A. Hermans, C. Ulens, S. Verstraeten, J. Helin, W. Depuydt, and M. De Paep for technical and administrative support.

## Author contributions

The study was designed by M.G., R.V. and L.M. Hardware design and motion capture recordings were done by L.M., A.B., R.V. and M.G. Data annotation, processing, animations and benchmarking were done by L.M. The macaque behavioral experiment was carried out by A.B. Eye fixation analysis was performed by L.M. and A.B. Statistical analysis was done by L.M. The article was written by L.M. with input from all authors.

