## Supplementary Information for "MacAction: Realistic 3D macaque body animation based on multi-camera markerless motion capture"

### Supplementary Figures

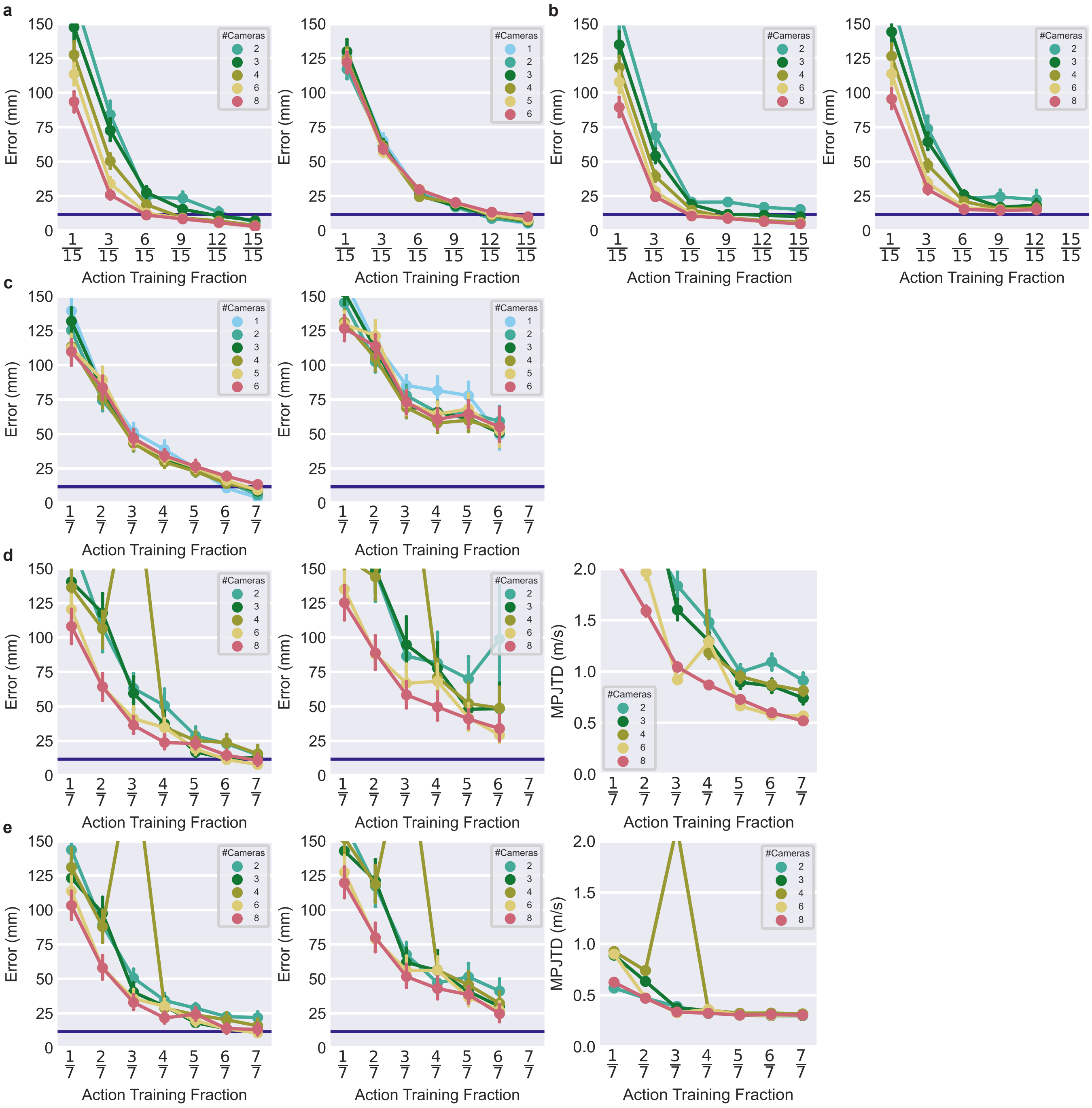


**Supplementary Figure 1: Errors for the submissive action training set and walking action performance.** **a,** Euclidean error for frames included in the fraction of the data that was used for training of DLC with RANSAC and structural refinement (left), vs. DANNCE (right). The horizontal line indicates the spatial voxel resolution (here $\sim$ 11.72mm) of the volumetric approach, DANNCE. In **a-e** points indicate the arithmetic means, and the vertical lines illustrate the 95% confidence intervals. **b,** Euclidean error for frames in the training (left) and test set (right) for spatiotemporally refined triangulation with Anipose. **c,** Euclidean error for frames in the training (left), and test set (right) for the walking action for DANNCE. **d,** Euclidean error for frames in the training (left), test set (middle), and MPJTD for the walking action with DLC and structurally refined triangulation (right). **e,** Euclidean error on frames in the training (left), test set (middle), and MPJTD (right) for the walking action with DLC and spatiotemporally refined triangulation.

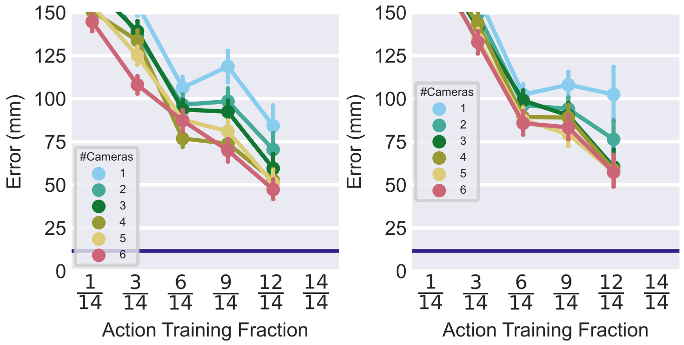


**Supplementary Figure 2: Multi-animal prediction with and without inclusion of other actions in the training.** Euclidean error for frames in the test set of the multi-animal action for DANNCE, with (left) and without (right) all other actions included in the training. The horizontal line indicates the spatial resolution (here, $\sim 11.72$mm) of the volumetric approach, DANNCE. Points indicate the arithmetic means, and the vertical lines illustrate the 95% confidence intervals.

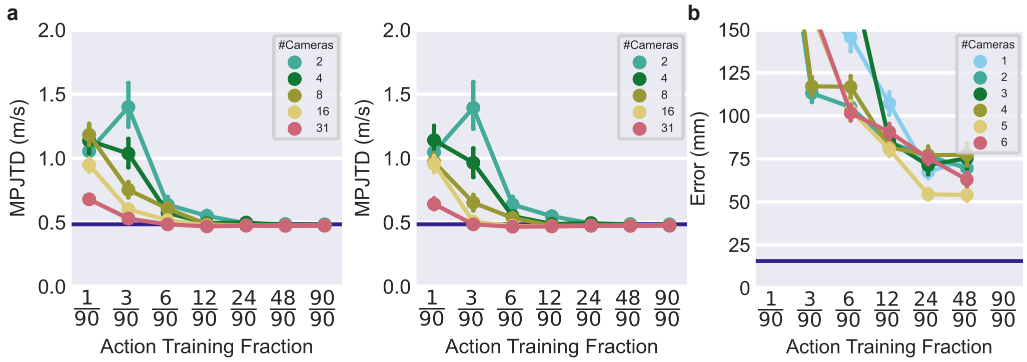


**Supplementary Figure 3: MPJTD for DLC without temporal refinement, and for the inclusion of other actions. a,** MPJTD for DLC using the direct linear transform (left) and additional structurally refined triangulation (right). The violet horizontal line indicates the ground-truth mean marker speed (0.49 m/s). In **a** and **b** the points indicate the arithmetic means, and the vertical lines illustrate the 95% confidence intervals. **b,** Euclidean error for frames included in the test set for DANNCE predicting the multi-agent interaction when including the dynamic single subject action in the training. The horizontal line in **b** indicates the spatial resolution (here, 15.625mm) of the volumetric approach, DANNCE.

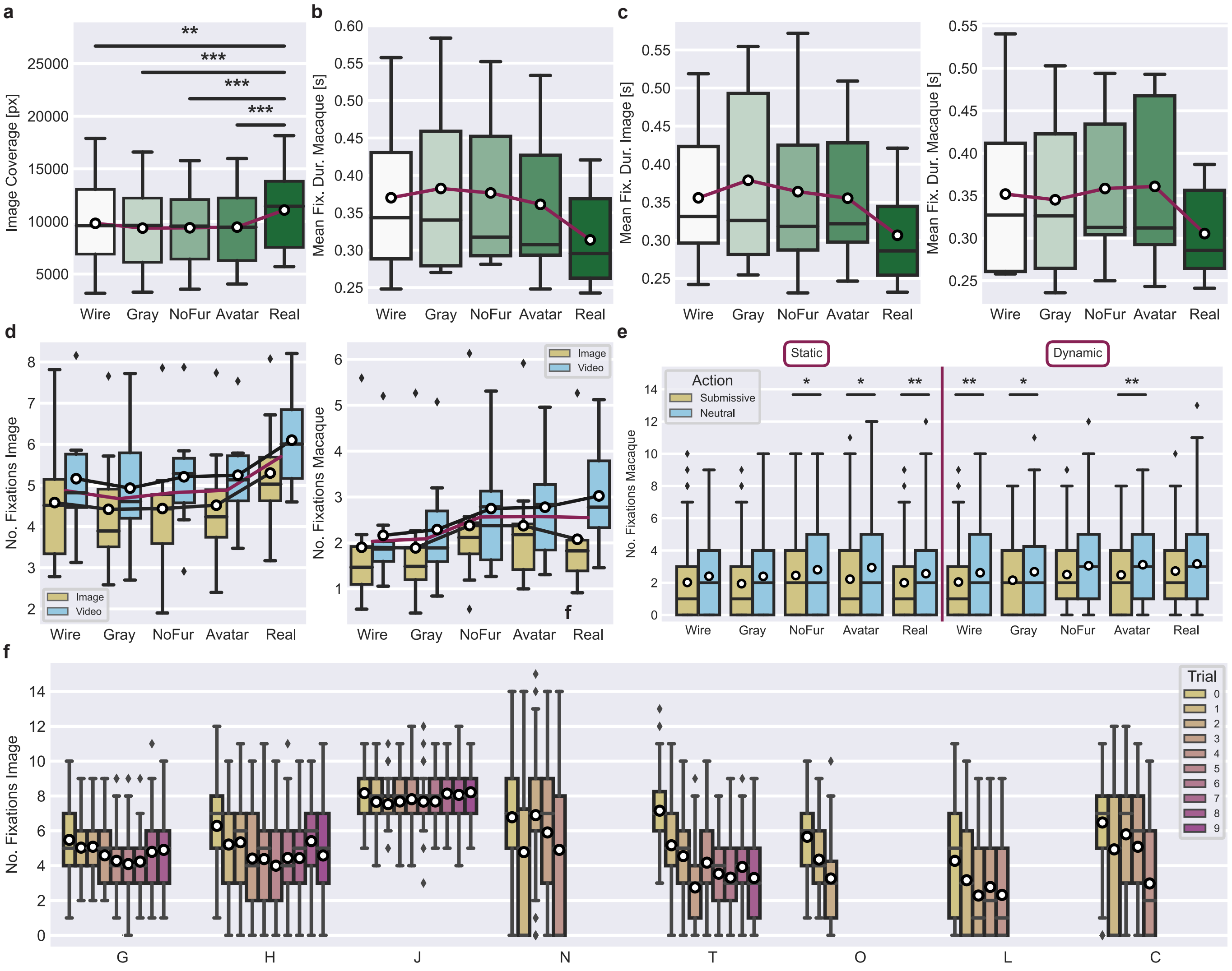


**Supplementary Figure 4: Extended results of the uncanny valley experiment.** **a,** The image coverage as function of realism including all trials. The plots in **c** show results of the first trials only, where all other panels include all trials of the individuals, respectively. The box plots show the median with interquartile range (IQR), whiskers at 1.5× the IQR and outliers depicted as diamonds. The arithmetic mean is shown as a white circle. In **a-d** arithmetic means are connected by a red line. Statistically significant pair-wise differences are indicated by horizontal lines accompanied with asterisks: $p^{**} < 0.01$ and $p^{***} < 0.001$. **b,** Box plot of mean fixation duration on the silhouette as function of render types. **c,** Box plot of the mean fixation duration on the image (left), and monkey silhouette (right) as function of render type, for the first trial only. **d,** The number of fixations on the image (left), and within the macaque silhouette (right) as function of render type, and split by presentation type. Arithmetic means per condition are connected by black lines, with the overall mean per render type indicated by a red line, omitting the additional white circles. **e,** Box plot showing the number of fixations of the macaque silhouette, split by action, render type, and presentation type. Image responses are shown on the left and fixation counts of videos on the right. **f,** Number of image fixations over trials and individuals.

### Supplementary Tables

**Supplementary Table 1: Related to Figure 4: Statistics for Image Coverage.** We conducted a one-way, repeated measures ANOVA to assess the effect of render type (realism) on the mean image coverage values from the first trials. The ANOVA revealed a significant influence of the render type: $F(4, 28) = 6.953, p = 0.005, \eta_g = 0.123$. Results of subsequently performed post-hoc pairwise, two-sided, t-tests with Benjamini-Hochberg correction are listed in the following table:

| | Group1 | Group2 | n1 | n2 | T-statistic | df | p-value | p-value adj. |
|---|---|---|---|---|---|---|---|---|
| Coverage | NoFur | Avatar | 8 | 8 | -0.8754 | 7 | 0.410 | 0.484 |
| Coverage | NoFur | Gray | 8 | 8 | 2.1644 | 7 | 0.067 | 0.096 |
| Coverage | NoFur | Real | 8 | 8 | -3.6559 | 7 | 0.008 | 0.041 |
| Coverage | NoFur | Wireframe | 8 | 8 | 0.0277 | 7 | 0.979 | 0.979 |
| Coverage | Avatar | Gray | 8 | 8 | 2.3751 | 7 | 0.049 | 0.082 |
| Coverage | Avatar | Real | 8 | 8 | -2.4313 | 7 | 0.045 | 0.082 |
| Coverage | Avatar | Wireframe | 8 | 8 | 0.8270 | 7 | 0.436 | 0.484 |
| Coverage | Gray | Real | 8 | 8 | -5.0378 | 7 | 0.002 | 0.015 |
| Coverage | Gray | Wireframe | 8 | 8 | -2.6997 | 7 | 0.031 | 0.077 |
| Coverage | Real | Wireframe | 8 | 8 | 2.7846 | 7 | 0.027 | 0.077 |

**Supplementary Table 2: Related to Figure 4: Negative Binomial Regression model for monkey fixation counts on all trials and experimental factors.** In order to analyze the monkey fixation counts as functions of Trial, Action, View, Presentation type (Video vs. Static), and Render type (Realism), we performed a binomial regression analysis. The corresponding model summary is shown in the following table:

| | Coefficient | 95%-confidence interval | Standarderror | *p*-value |
|---|---|---|---|---|
| Intercept | 1.16 | from 2.91 to 3.50 | 0.048 | $< 0.0001$ |
| ActionWalk | 0.22 | from 1.20 to 1.30 | 0.02 | $< 0.0001$ |
| ViewC | -0.023 | from 0.91 to 1.05 | 0.036 | 0.53 |
| ViewD | -0.13 | from 0.82 to 0.94 | 0.036 | 0.0004 |
| ViewE | -0.30 | from 0.69 to 0.80 | 0.037 | $< 0.0001$ |
| ViewF | -0.25 | from 0.73 to 0.84 | 0.037 | $< 0.0001$ |
| ViewG | -0.18 | from 0.78 to 0.90 | 0.036 | $< 0.0001$ |
| ViewH | -0.31 | from 0.68 to 0.79 | 0.037 | $< 0.0001$ |
| VideoStatic | -0.15 | from 0.82 to 0.89 | 0.02 | $< 0.0001$ |
| IndividualGin | -0.024 | from 0.90 to 1.06 | 0.043 | 0.57 |
| IndividualHyde | 0.11 | from 1.03 to 1.21 | 0.042 | 0.011 |
| IndividualJekyll | 1.05 | from 2.65 to 3.10 | 0.039 | $< 0.0001$ |
| IndividualLibre | -0.73 | from 0.43 to 0.53 | 0.053 | $< 0.0001$ |
| IndividualNacho | 0.05 | from 0.96 to 1.15 | 0.046 | 0.28 |
| IndividualOdin | -0.11 | from 0.80 to 0.99 | 0.054 | 0.039 |
| IndividualTonic | -0.51 | from 0.55 to 0.66 | 0.045 | $< 0.0001$ |
| RenderAvatar | 0.0014 | from 0.94 to 1.06 | 0.031 | 0.96 |
| RenderGray | -0.18 | from 0.79 to 0.89 | 0.032 | $< 0.0001$ |
| RenderReal | -0.02 | from 0.92 to 1.04 | 0.031 | 0.51 |
| RenderWireframe | -0.19 | from 0.77 to 0.88 | 0.032 | $< 0.0001$ |
| Trial | -0.07 | from 0.93 to 0.94 | 0.0041 | $< 0.0001$ |

**Supplementary Table 3: Related to Figure 4: Interaction between Render type and stimulus dynamics.** Likelihood-ratio tests (LRTs) for the *image* fixation counts for the first trial. The factors Render and Video (Presentation type) were each significant for modeling, unlike their interaction.

| | Model | Criterion variable | Predictor variables | df-model | df | $\chi^2$-statistic | p-value |
|---|---|---|---|---|---|---|---|
| LRT 1 | $\Theta_0$ | ImageFixations | Action + View + Individual | 16 | | | |
| | $\Theta$ | ImageFixations | Action + View + Individual + Video | 17 | | | |
| | | | | | 1 | 41.01 | $< 0.0001$ |
| LRT 2 | $\Theta_0$ | ImageFixations | Action + View + Individual | 16 | | | |
| | $\Theta$ | ImageFixations | Action + View + Individual + Render | 20 | | | |
| | | | | | 4 | 35.66 | $< 0.0001$ |
| LRT 3 | $\Theta_0$ | ImageFixations | Action + View + Individual + Video + Render | 21 | | | |
| | $\Theta$ | ImageFixations | Action + View + Individual + Video + Render + Render * Video | 25 | | | |
| | | | | | 4 | 3.05 | 0.5502 |

**Supplementary Table 4: Related to Figure 4: Interaction between Render type and stimulus dynamics.** Likelihood-ratio tests (LRTs) for fixation counts on *monkey silhouettes* for the first trial. The influences of the factors Render and Video (Presentation type) were significant, unlike their interaction.

| | Model | Criterion variable | Predictor variables | df-model | df | $\chi^2$-statistic | p-value |
|---|---|---|---|---|---|---|---|
| LRT 1 | $\Theta_0$ | MonkeyFixations | Action + View + Individual | 16 | | | |
| | $\Theta$ | MonkeyFixations | Action + View + Individual + Video | 17 | | | |
| | | | | | 1 | 57.07 | < 0.0001 |
| LRT 2 | $\Theta_0$ | MonkeyFixations | Action + View + Individual | 16 | | | |
| | $\Theta$ | MonkeyFixations | Action + View + Individual + Render | 20 | | | |
| | | | | | 4 | 34.04 | < 0.0001 |
| LRT 3 | $\Theta_0$ | MonkeyFixations | Action + View + Individual + Video + Render | 21 | | | |
| | $\Theta$ | MonkeyFixations | Action + View + Individual + Video + Render + Render * Video | 25 | | | |
| | | | | | 4 | 1.60 | 0.8086 |

**Supplementary Table 5: Related to Figure 4: Two-sided paired Wilcoxon tests on combinations of the factors Render and Video (Presentation type) for action discrimination including all trials.** In single-pair comparisons for different conditions, mean differences across action types were significantly higher in more than half of the cases.

| | Subset | Group1 | Group2 | W-statistic | n | p-val | CLES |
|---|---|---|---|---|---|---|---|
| 0 | Wireframe+Static | Submissive | Neutral | 4.0000 | 8 | 0.0547 | 0.3906 |
| 1 | Gray+Static | Submissive | Neutral | 5.0000 | 8 | 0.0781 | 0.4297 |
| 2 | NoFur+Static | Submissive | Neutral | 1.0000 | 8 | 0.0156 | 0.3438 |
| 3 | Avatar+Static | Submissive | Neutral | 1.0000 | 8 | 0.0156 | 0.3438 |
| 4 | Real+Static | Submissive | Neutral | 0.0000 | 8 | 0.0078 | 0.2656 |
| 5 | Wireframe+Dynamic | Submissive | Neutral | 0.0000 | 8 | 0.0078 | 0.2344 |
| 6 | Gray+Dynamic | Submissive | Neutral | 0.0000 | 7 | 0.0225 | 0.3516 |
| 7 | NoFur+Dynamic | Submissive | Neutral | 4.0000 | 8 | 0.0547 | 0.3750 |
| 8 | Avatar+Dynamic | Submissive | Neutral | 0.0000 | 8 | 0.0078 | 0.3281 |
| 9 | Real+Dynamic | Submissive | Neutral | 3.0000 | 7 | 0.0759 | 0.4297 |

**Supplementary Table 6: Related to Figure 4: Interaction between the factors Render and Action.** Likelihood-ratio tests (LRTs) for fixation counts on *monkey silhouettes* for the first trial, comparing real footage and avatar only. There was a significant contribution of the Action type to modeling fixation counts. However, the Render type and its interaction with the Action type did not significantly contribute to the model, demonstrating the similar perception of avatar and real videos in subjects.

| | Model | Criterion variable | Predictor variables | df-model | df | $\chi^2$-statistic | p-value |
|---|---|---|---|---|---|---|---|
| LRT 1 | $\Theta_0$ | MonkeyFixations | View + Individual + Video | 16 | | | |
| | $\Theta$ | MonkeyFixations | View + Individual + Video + Render | 17 | | | |
| | | | | | 1 | 0.29 | 0.5885 |
| LRT 2 | $\Theta_0$ | MonkeyFixations | View + Individual + Video | 16 | | | |
| | $\Theta$ | MonkeyFixations | View + Individual + Video + Action | 17 | | | |
| | | | | | 1 | 23.43 | < 0.0001 |
| LRT 3 | $\Theta_0$ | MonkeyFixations | View + Individual + Video + Render + Action | 18 | | | |
| | $\Theta$ | MonkeyFixations | View + Individual + Video + Render + Action + Render * Action | 19 | | | |
| | | | | | 1 | 0.46 | 0.498 |

### Supplementary Note

#### Re-identification algorithm

The following algorithm re-associates the 2D markers with the corresponding $N_A$ animal identities for all $N_M$ hierarchical keypoints in an action sequence of length $T$. We denote this mapping across $N_C$ camera views in the following $D^2$, and it is given by a tensor with dimensionality $N_C \times N_M \times N_A \times T$. Ideally starting from a labeled keyframe without another tracked animal closeby, the animal identities of the root markers are arbitrarily initialized. Then, by iterating along the skeletal hierarchy, the keypoints are associated with their parent node. We denote this association for a given time step $t$ and keypoint $k$ as $I^3_{k,t}$, a vector of size $N_A$. The function *Associate3DHierarchy* exclusively assigns plausible 3D markers $M^3_{k,t} \in \mathbb{R}^{3\times N_A}$ by minimal Euclidean distance. Note that $M^3_{k,t}$ is derived from distinct camera sets. After the first time step, the flag $kin$ changes, triggering the association of animal identities to the preceding time step by the routine $Associate3DTime$, which also links markers by Euclidean distance. Unless there is a significant increase in 3D Euclidean distance $d_t$ that exceeds a constant $c$, the association of the $N_A$ markers of the k-th keypoint remains the same. Otherwise, animal identities are resolved by their parental markers, with the association changing back to over time for subsequent iterations. Resolved associations in 3D are then updated in $I^3$ after each time step and keypoint. Finally, these animal identities are projected into each camera view, re-identifying each marker in the image with its corresponding animal.

**Algorithm 1:** Algorithm for 3D individual identification

```
Result: Disambiguated animal identities for 2D markers
kin ← False
M^2 ← Sort tracked 2D markers by skeleton hierarchy
I^3 ← Initialize 3D individual association
for t ∈ {1, ..., T} do
    for k ∈ {1, ..., N_M} do
        I^3_{k,t}, M^3_{k,t} ← Initialize3D(M^2)
        if kin is True then
            d_t, I^3_{k,t} ← Associate3DTime(M^3_{k,t}, M^3_{k,t-1}) // Associate with last time step
            if d_t ≥ c then
                kin ← False
            end
        end
        if k > 1 and kin is False then
            I^3_{k,t} ← Associate3DHierarchy(M^3_{k,t}, M^3_{k-1,t}) // Associate with parental keypoint
            if t > 1 then
                kin ← True
            end
        end
        I^3 ← Update(I^3_{k,t})
    end
end
D^2 ← ProjectIdentities(I^3)
```

##### Posing single joints in the avatar model

We distinguish between joints with two and three rotational degrees of freedom (DOF). For two DOF, consider the column vector $a = (a_1, a_2, a_3)^T$ as the relative position of the succeeding element in the avatar's kinematic tree. We denote $b = (b_1, b_2, b_3)^T$ as the respective tracked version of $a$ in the same coordinate system. After transforming both vectors into spherical coordinates $(r_x, \theta_x, \varphi_x)$, we subtract their angles to form $\varphi_{diff} = \varphi_b - \varphi_a$ and $\theta_{diff} = \theta_b - \theta_a$. We construct a rotation matrix around the z-axis with $\varphi_{diff}$ and rotate $a$ with it. With this transformed vector and b, we construct the orthogonal vector $b \times aR_z(\varphi_{diff})$ by cross-product, norm it and denote it as $e$. To transform $e$ by $\theta_{diff}$, we use the matrix form of an axis-angle rotation. In general, the counter-clockwise rotation around an unit vector $u = (u_1, u_2, u_3)^T$ by an angle $\alpha$ is given by [1]:

$$R_u(\theta) = \begin{bmatrix} q + u_1^2(1-q) & u_1u_2(1-q) - u_3p & u_1u_3(1-q) + u_2p \\ u_2u_1(1-q) + u_3p & q + u_2^2(1-q) & u_2u_3(1-q) - u_1p \\ u_3u_1(1-q) - u_2p & u_3u_2(1-q) + u_1p & q + u_3^2(1-q) \end{bmatrix}$$
$$\text{with} \quad p = \sin\theta \quad \text{and} \quad q = \cos\theta.$$

So that the transformation of $a$ to $b$ can be written as $R_1 = R_e(\theta_{diff})R_z(\varphi_{diff})$. We adjust the sign of $\theta_{diff}$ according to the orientation of $e$ to rotate in the appropriate direction. This technique establishes the pose for joints with two DOFs, keeping the roll of the transformed $a$ intact. The two transformations relate to a yaw and pitch transformation of the animated vector.

For three-DOF joints, each vector — both animated and tracked — relates to a secondary vector required to define the roll. Here, $c$ and $d$ denote the secondary vectors for animation and tracking, respectively. Note that, the rotations between $a$, $c$ and $b$, $d$ are generally not the same. Thus, we transform $c$ by $R_1$ and determine the roll that makes $R_b(\gamma)R_1c$ coplanar with $b$ and $d$:

$$A = \left[\begin{array}{c|c|c} b & d & R_b(\gamma)R_1c \end{array}\right], \text{and}$$
$$\det A \stackrel{!}{=} 0,$$

where $A$ denotes a column matrix composed of these three column vectors. The angles that satisfy this condition are given by:

$$\gamma = \pi n - \arctan(\frac{x}{y})$$
$$\text{with} \quad y \neq 0 \quad \text{and} \quad n \in \mathbb{Z},$$

where $x$ and $y$ can be expressed as:

$$x = (d_2c_3 - d_3c_2)b_1 + (d_3c_1 - d_1c_3)b_2 + (d_1c_2 - d_2c_1)b_3,$$
$$y = (-d_2c_1 - d_1c_2)b_1b_2 + (-d_1c_3 - d_3c_1)b_1b_3 + (-d_3c_2 - d_2c_3)b_2b_3$$
$$+(d_2c_2 + d_3c_3)b_1^2 + (d_1c_1 + d_3c_3)b_2^2 + (d_1c_1 + d_2c_2)b_3^2.$$

We determine whether we need to add $\pi$ to the angle by scalar product with $d$ and the transformed $c$. All other solutions represent the same two rotations.
